# Label-Free Identification of Human Eosinophils Using 808 nm Side Scatter

**DOI:** 10.64898/2026.06.04.730064

**Authors:** Krittika Ralhan, Fanuel Messagio, Joost M. Lambooij, T. Tak

**Affiliations:** Beckman Coulter Life Sciences; Leiden University Center of Infectious Diseases, Leiden University Medical Center, Leiden, The Netherlands; Flow Cytometry Core Facility, Leiden University Medical Center, Leiden, The Netherlands

## Abstract

Accurate identification and quantification of eosinophils is critical for the diagnosis and monitoring of eosinophil-associated disorders. While flow cytometry remains a powerful tool for leukocyte characterization, conventional instruments equipped with 405 nm or 488 nm side scatter (SSC) detectors offer limited resolution for eosinophil discrimination overlap in scatter with neutrophils. Using a spectral flow cytometer equipped with six distinct SSC detectors, we report a novel, label-free approach for eosinophil detection leveraging high 808 nm near-infrared SSC (IRSSC) uniquely observed in human eosinophils. This optical signature is independent of antibody labeling, activation fixation, or permeabilization, and shows strong concordance with conventional CD66b/CD16 gating strategies (R = 0.997). Notably, the high 808 nm SSC is absent in murine eosinophils, suggesting a species-specific structural feature such as in human eosinophils. These findings establish IRSSC as a robust, reagent-free biomarker for eosinophil detection, with broad implications for both clinical diagnostics and translational immunology.

## Introduction

Eosinophils are implicated in type 2 immune responses and the pathogenesis of diverse diseases including asthma, parasitic infections, and eosinophilic gastrointestinal disorders (1, 2, 3). Their identification and enumeration in whole blood or tissue preparations are critical for diagnosis and monitoring of eosinophilic conditions.

Most flow cytometry platforms utilize light scatter properties at 405 and 488 nm. On these SSC detectors, eosinophils have a high SSC compared to lymphocytes, monocytes and neutrophils (4, 5, 6). There is an overlap, however, between the neutrophil and eosinophil populations, which prevents the identification of eosinophils using scatter alone. Immunophenotypic discrimination using CD16 and CD66b further necessitates staining protocols, thereby increasing complexity and potentially altering native cellular features (7, 8).

Label-free methods to identify eosinophils using scatter do exist. For example, depolarization of 488 nm SSC is shown to uniquely identify eosinophils (9). However, the use of fibre-optics in many modern flow cytometers precludes the detection of depolarization (10). In addition, eosinophils are known to have a unique autofluorescence (11)) and they can be identified using image cytometry (12).

The CytoFLEX mosaic Spectral Detection Module extends SSC detection to six lasers, ranging from 355 nm to 808 nm, enabling additional characterization of cells. In this study, we evaluate the capability of different SSC wavelengths to resolve leukocyte populations in a label-free manner. Our research shows that eosinophils are uniquely displaying a high SSC signal on high wavelength lasers. This highlights the structural uniqueness of human eosinophils and introduces a method with potential for rapid, label-free flow cytometric quantification of eosinophils.

## Materials and Methods

### Sample Preparation

Human peripheral blood from healthy human volunteers was obtained from the LUMC Volunteer Donor Service (LuVDS, application number LuVDS25.010) under institutional ethics approval. Erythrocytes were removed via isotonic ammonium chloride-based lysis (RBC Lysis Buffer, BioLegend) according to manufacturers’ protocols. Leukocytes were stained with fluorochrome-conjugated antibodies (Table 1) in the presence of TruStain FcX (BioLegend), True-Stain Monocyte Blocker (BioLegend) and Brilliant Stain Buffer Plus (BD Biosciences) according to manufacturers’ protocols in PBS supplemented with 1% Fetal Bovine Serum (FACS Buffer). Fixation and permeabilization were performed using the eBioscience Foxp3 /Transcription Factor Staining Buffer Set (Thermo Fisher Scientific) according to manufacturers’ instruction, which was analyzed directly or followed by 90% ice-cold Methanol fixation (13), when indicated.

**Table 1.**
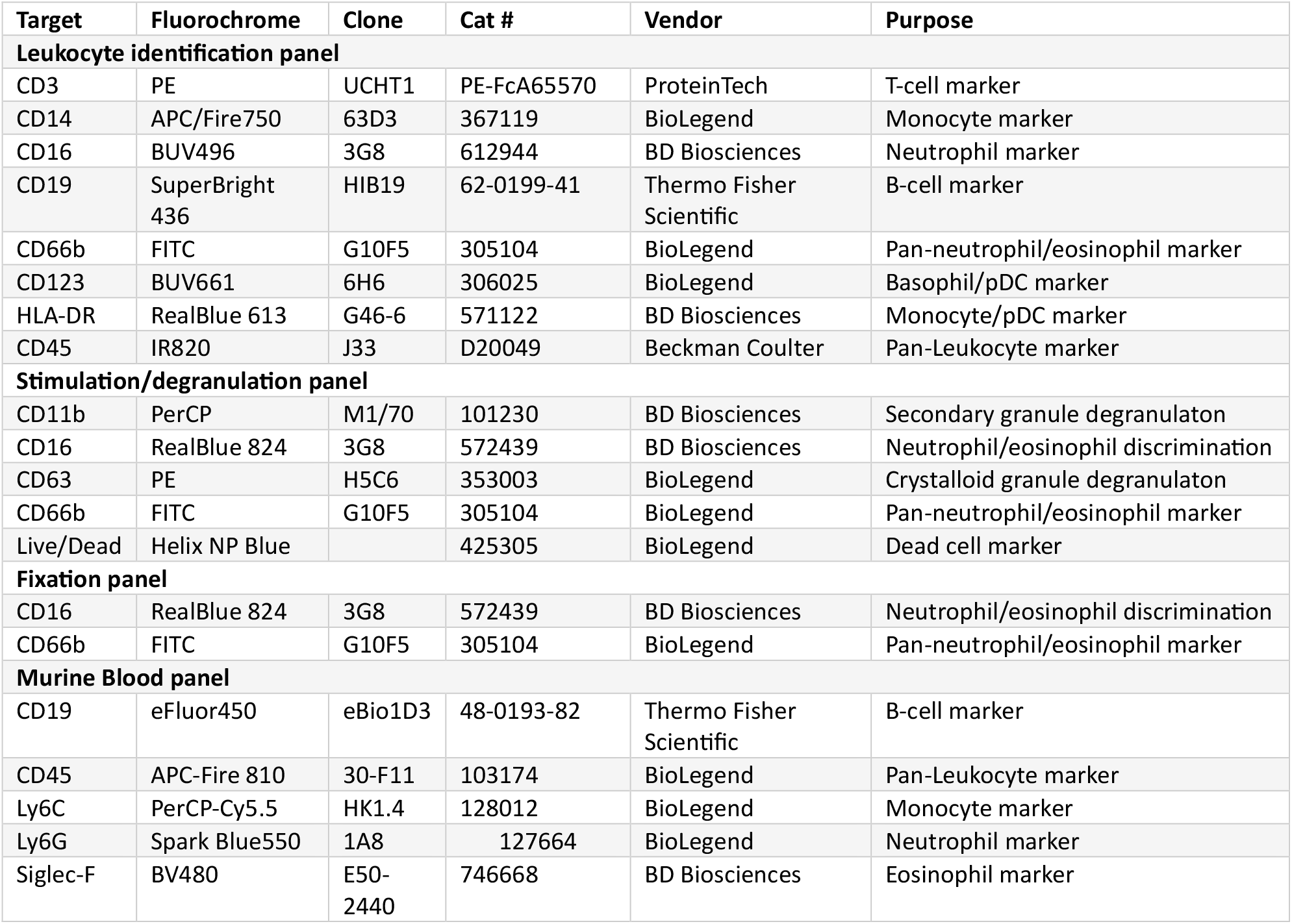
Flow Cytometry panels:

### Sample Stimulation

To induce eosinophil degranulation. RBC lysed WBC were stimulated with PMA (100 ng/ml), Ionomycin (1µg/ml) or PMA with ionomycin in 100µl RPMI1640 supplemented with 8% FBS for 15 minutes at 37°C. The reaction was stopped by adding 100µl ice-cold FACS buffer. Samples were stained with fluorescently labelled antibodies (table 1) in the presence of TruStain FcX and True-stain monocyte blocker and washed twice before analysis.

Murine blood samples were obtained in accordance with the Guide for the Care and Use of Laboratory Animals of Laboratory Animal Research, and with approval from the Dutch Central Authority for Scientific Procedures on Animals (CCD; license number: AVD1160020174364). Blood was obtained from C57BL/6 mice and stained according to the procedures described in OMIP-104 14).

### Flow Cytometry

Blood samples were analyzed using a CytoFLEX LX Flow Cytometer equipped with a mosaic 88 Spectral Detection Module (Beckman Coulter Life Sciences). The flow cytometer offers six SSC detectors, one for each laser: 355, 405, 488, 561, 638, and 808 nm (Supplemental table 1 and Supplemental Figure 1). Samples were acquired using standardized assay settings at a flow rate of 60ul/min with at least 100,000 leukocytes detected per sample. Unmixing was performed using poisson-hybrid unmixing with CytExpert V1.2.0.4 taking into account two autofluorescent cell populations (monocytes and eosinophils).

### Data analysis

Cell populations in human peripheral blood samples were identified as 405 nm SSC (VSSC)^high^/CD66b^+^/CD14^-^/CD16^+^ neutrophils, VSSC^high^/CD66b^+^/CD14^-^/CD16^-^ eosinophils, VSSC^low^/CD66b^-^/CD3^+^ T cells, VSSC^low^/CD66b^-^/CD19^+^ B cells, VSSC^low^/CD66b^-^/CD3^-^/CD19^-^ /CD123^+^/HLA-DR^-^ basophils, VSSC^low^/CD66b^-^/CD3^-^/CD19^-^/CD123^+^/HLA-DR^+^ pDC and VSSC^low^/CD66b^-^/CD3^-^/CD19^-^/CD123^-^/HLA-DR^+^/CD14^+/dim^/CD16^+/-^ monocytes.

Cell populations in murine blood were identified as Live/CD45^+^/Ly6G^+^/Siglec-F^-^ neutrophils, Live/CD45^+^/Ly6G^-^/Siglec-F^+^ eosinophils, Live/CD45^+^/Ly6G^-^/Siglec-F^-^/Ly6C^+^/CD19^-^ monocytes and Live/CD45^+^/Ly6G^-^/Siglec-F^-^/Ly6C^-^/CD19^+^ B-cells.

### Data Analysis

Samples were analysed and visualized using OMIQ (Dotmatics). Graphs were created using GraphPad Prism 10.2.3 (Dotmatics). For visualization, fluorescence data was arcsinh transformed with cofactor 6000.

To provide a measurement that allows comparisons of measurements on different SSC detectors, an SSC ratio was calculated for each cell population by dividing the Scatter intensity by the scatter intensity of CD3+ T-cells, as this was the population with the lowest SSC on all channels.

## Results

### Label-free detection of eosinophils using 808nm SSC

To assess the resolving capability of each SSC detector, side scatter data were plotted against FSC-A for each detection wavelength (Figure 2A). As the laser wavelength increased, the separation between neutrophils and eosinophils became progressively more pronounced, while the separation between other populations decreased.

**Figure 1.**
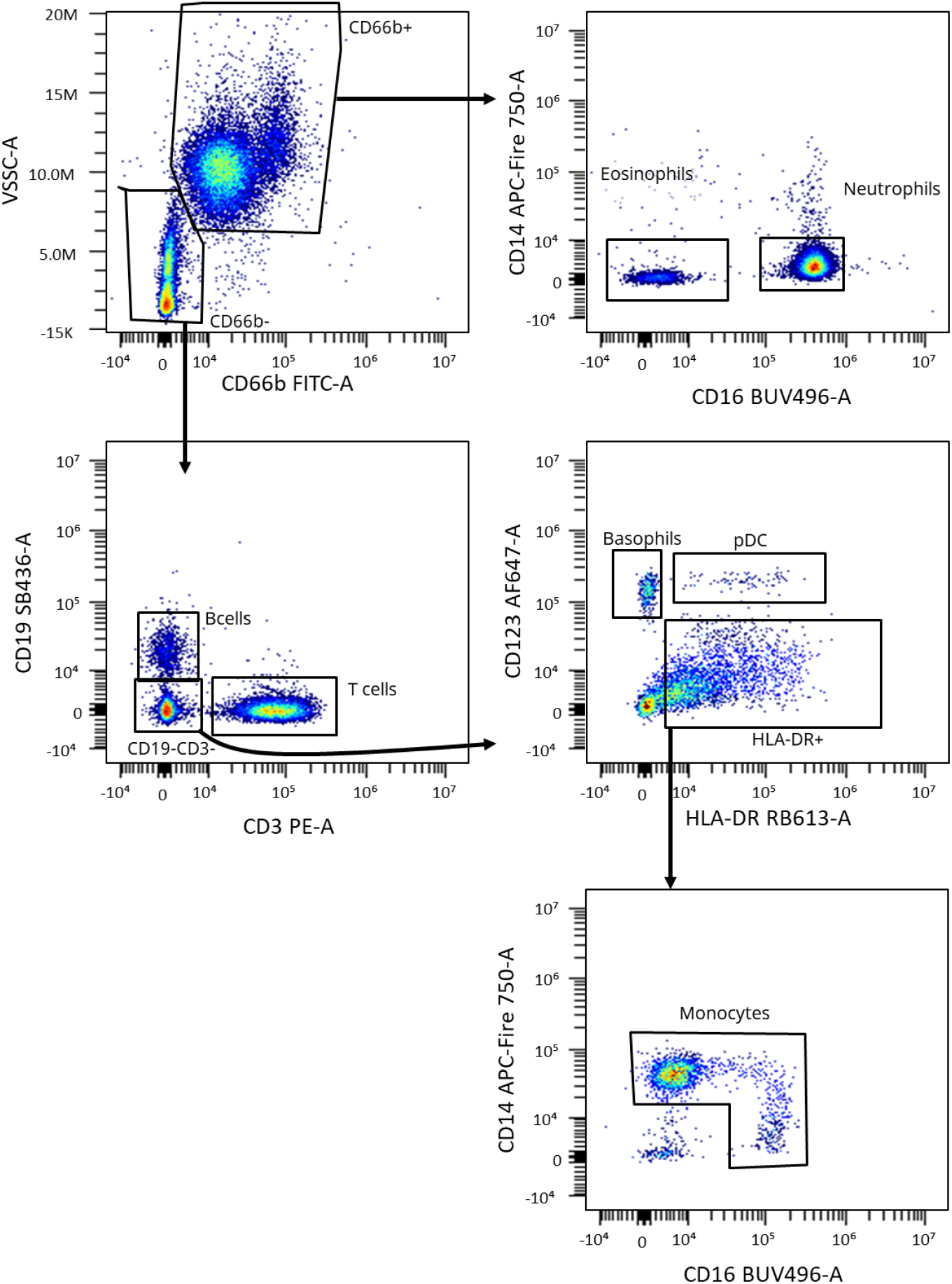
Gating strategy. RBC lysed Human whole blood samples were identified using the gating strategy shown. Representative figure of 5 samples.

**Figure 2.**
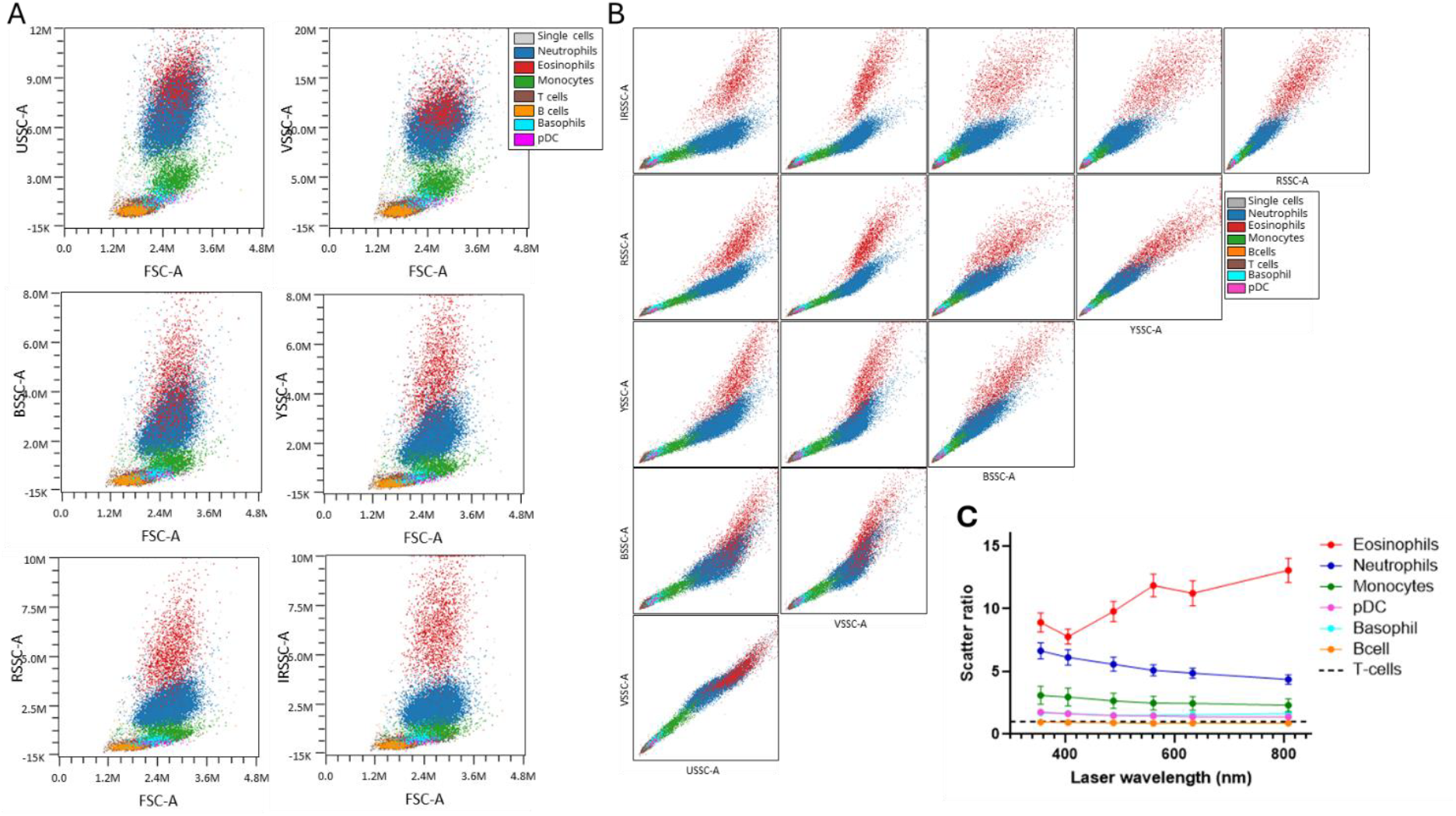
Overlays of different cell populations were plotted as FSC-A versus the different SSCs (A) show an increase in the separation between neutrophils and eosinophils with increasing scatter wavelength. Scatter NxN plots (B) show that especially IRSSC-A plotted against VSSC-A allows label-free identification of eosinophils. The scatter ratio reveals that only eosinophils display an increased scatter with an increasing wavelength. Plots represent a representative example of at least 3 experiments (A,B) or means +/-SD of 3-5 samples.

To determine which combination of scatters is best to resolve eosinophils from other cell types and determine whether any other cell types could be discerned using scatter alone, an NxN plot was created for the different SSCs. The NxN plot showed that eosinophils are best identified using 808 nm Infra-red (IR) SSC plotted against 405nm Violet (V) SSC, although any combination of 638 Red (R))SSC or IRSSC plotted against 355nm UVSSC or VSSC allows label-free identification of eosinophils. Besides eosinophils, none of the other cell populations investigated could be identified using a combination of SSCs. Interestingly, murine eosinophils did not show the same separation from other cell populations (supplementary figure 2).

### Relative quantification of SSC

To quantitatively compare differences in scatter between cell populations irrespective of the absolute SSC for each detector, an SSC ratio was calculated. This revealed that the increase in SSC with wavelength was specific to eosinophils, while the relative separation of monocytes, pDC, basophils and neutrophils decreased with an increasing SSC wavelength.

### Resistance of IRSSC signal irrespective of sample treatment

The high IRSSC signal from eosinophils was shown to be independent of labeling procedure, as unstained samples showed a similar IRSSC^high^ population to antibody-stained samples (Figure 3A). Furthermore, the high IRSSC signal was resistant to cell fixation and permeabilization, since either fixation and permeabilization or methanol fixation did not reduce the IRSSC signal (Figure 3B).

**Figure 3.**
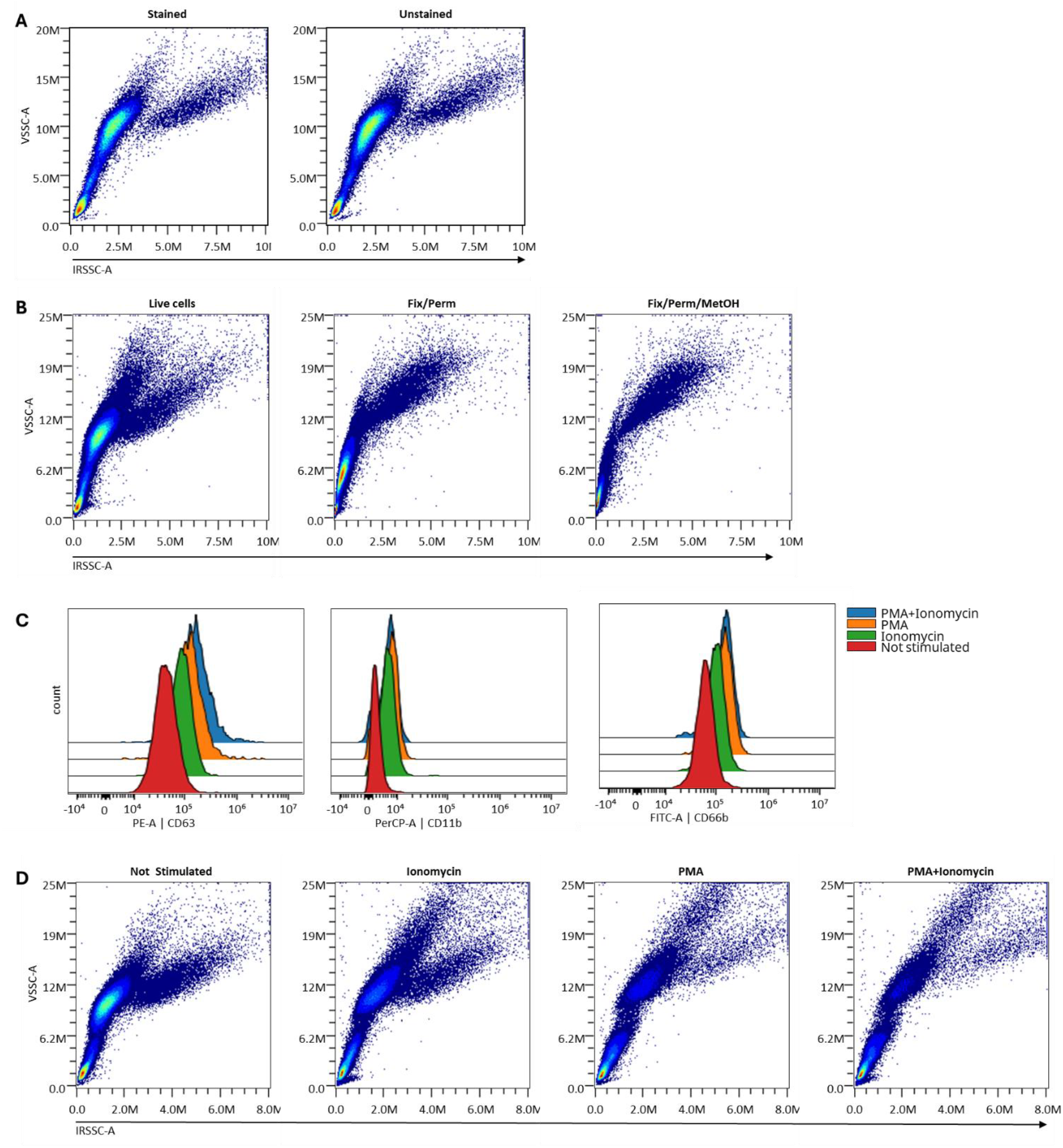
Eosinophil-specific IR-SSC is independent of sample treatment. Eosinophils display an increased IRSSC independent of antibody staining (A) or fixation/permeabilization (B). Eosinophil degranulation was quantified by upregulation of degranulation markers CD11b, CD66b and CD63 (C). Cellular activation did not affect eosinophil IRSSC signal (D). Data are concatenated data obtained from 3 experiments.

**Figure 4.**
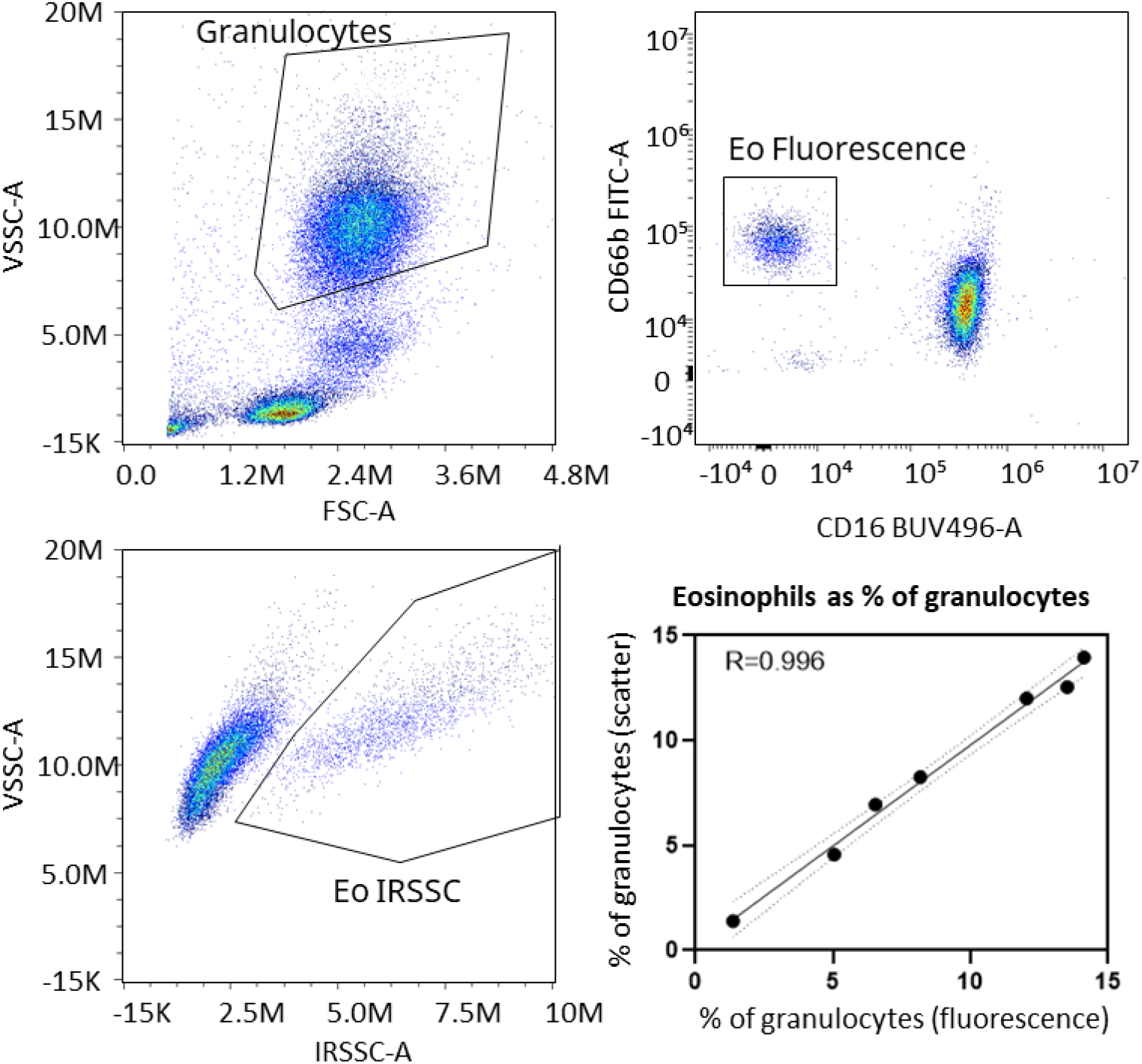
Comparison of quantification methods. Eosinophils were identified from the granulocyte population (top left) using either CD66b and CD16 expression (top right) or V-SSC and IR-SSC intensities (lower left). The two resulting percentages of granulocytes were plotted in a correlogram (lower right). Solid line indicates the linear fit, dashed lines indicate 95% confidence interval of the fit.

Stimulation of RBC lysed whole blood with PMA and/or Ionomycin induced a strong cell activation and degranulation phenotype as determined by increased expression of activation and degranulation markers CD11b, CD66b and CD63 (Figure 3C). However, there was no difference in IRSSC signal between conditions (Figure 3D), indicating that the high IRSSC signal is retained upon eosinophil degranulation.

### Comparison of quantification methods

The proportion of eosinophils within the granulocyte population was quantified using two independent strategies: CD16/CD66b immunophenotyping and side-scatter–based separation (VSSC/IRSSC). Both methods produced highly concordant results, with a Pierson correlation coefficient of 0.996 (N=7), indicating near-perfect agreement between marker-based and scatter-based eosinophil identification. This demonstrates that eosinophils can be reliably quantified using either approach under the applied experimental conditions.

## Discussion

Our findings demonstrate the use of high-wavelength SSC, specifically 808 nm, as a label-free method for eosinophil detection in human blood. The granule-rich morphology of eosinophils contributes to distinctive optical scattering behavior. Eosinophils are characterized by high side-scatter intensity and depolarized light scatter properties compared with other leukocyte populations, owing to their electron-dense crystalline granules. (figure 2C). (5)The high SSC intensity at high wavelengths is likely to originate from structures unique to eosinophils, such as eosinophil crystalloid granules. These granules contain a core of hexagonal bipyramidal Charcot-Leyden crystals (CLC) that have a relatively high refractive index (RI)(15, 16). The wavelength-dependent effect is consistent with Mie scattering theory, which predicts that scattering efficiency and angular distribution are influenced by both the size and refractive index of intracellular structures relative to the wavelength of incident light. (17, 18, 19) The large, crystalline, and optically dense granules of eosinophils maintain strong scattering even at near-infrared wavelengths, whereas the finer and less refractile granules of neutrophils and monocytes scatter less efficiently as wavelength increases (20, 21, 22)). Consequently, eosinophils may become optically more distinct at longer wavelengths, supporting the use of multi-wavelength SSC detection to improve granulocyte subclass resolution.

Interestingly, galectin-10, the main component of CLC is also expressed in basophils and basophils have also been shown to contain CLC, yet do not display a high IR-SSC intensity. (23) Differences in granule size or CLC content between eosinophils and basophils may explain the difference between these cell types. Indeed, prior work on polarized light scattering has shown that light of different wavelengths (488-630nm) scattered by eosinophils is depolarized. (24) Theoretical modeling of granule scattering demonstrated that the depolarization of light by eosinophils could be explained solely by the size of their granules. (25) Similarly, murine eosinophils did not display a high IR-SSC. This difference likely arises from interspecies variation in granule morphology and refractive index. Human eosinophils contain major basic protein– rich crystalline cores that strongly scatter light, whereas murine eosinophil granules are smaller, less crystalline, and exhibit reduced refractive contrast with the cytoplasm (26).

Activation of eosinophils with PMA and Ionomycin induced degranulation of crystalloid granules, as shown by an upregulation of CD63 (Figure 3C). Despite the observed degranulation, the intensity of IR-SSC did not decrease with degranulation. Even stimulation with strong stimuli does not fully degranulate eosinophils, as demonstrated by a similar lack of decrease in V-SSC intensity. Thus, IRSSC is likely suitable for detection of eosinophils in wide variety of conditions.

The strong correlation between CD16/CD66b gating and VSSC/IRSSC discrimination suggests that eosinophils can be robustly identified based on their distinct optical properties, independent of surface marker expression. These distinct optical properties are independent of both cellular activation and common sample fixation methods. The clinical and translational implications of this approach are clear. Label-free SSC detection enables high-throughput, cost-effective eosinophil enumeration without the need for antibody reagents or extensive sample processing. This technique could be readily implemented in diagnostic laboratories, and resource-limited settings, and offers potential for integration into automated cytometry workflows for hematological profiling.

## Acknowledgements

We would like to thank the FCF operators and students from the LUMC Flow cytometry Core Facility for their work validating and maintaining the CytFLEX LX and mosaic 88 flow cytometer.

## Notes

### Competing Interest Statement

KR and FM are employees of Beckman Coulter, the manufacturer of the CytoFLEX LX and mosaic88. TT and JL declare no competing interests

