## Supplemental table and figures for "Label-Free Identification of Human Eosinophils Using 808 nm Side Scatter"

Supplemental table 1: Optical layout of mosaic88

| 355 nm | 405nm | 488nm | 561nm | 638nm | 808nm |
|---|---|---|---|---|---|
| *355/8* | *405/8* | | | | |
| 372/15 | 426/12 | | | | |
| 387/15 | 440/16 | | | | |
| 427/15 | 455/15 | | | | |
| 443/16 | 472/18 | *488/8* | | | |
| 458/15 | 508/20 | 508/20 | | | |
| 473/15 | 525/15 | 525/15 | | | |
| 513/30 | 541/17 | 541/17 | | | |
| 541/26 | 579/22 | 579/22 | *561/8* | | |
| 582/29 | 599/18 | 599/18 | 577/20 | | |
| 612/31 | 616/17 | 616/17 | 597/21 | *638/12* | |
| 659/20 | 659/20 | 659/20 | 616/17 | 659/20 | |
| 678/18 | 678/18 | 678/18 | 659/20 | 678/18 | |
| 696/19 | 696/19 | 696/19 | 678/18 | 696/19 | |
| 716/20 | 717/20 | 716/20 | 696/19 | 716/20 | |
| 738/22 | 738/22 | 738/22 | 720/27 | 738/22 | |
| 760/23 | 760/23 | 760/23 | 748/30 | 760/23 | |
| 783/23 | 783/23 | 783/23 | 778/29 | 783/23 | *808/12* |
| 840/20 | 840/20 | 840/20 | 840/20 | 840/20 | 840/20 |
| 877/55 | 877/55 | 877/55 | 877/55 | 877/55 | 877/55 |
| 927/45 | 927/45 | 927/45 | 927/45 | 927/45 | 927/45 |

Optical layout, depicted as wavelength/bandpass. *Italicized* filters indicate SSC filters for SSC detection

Supplemental figures

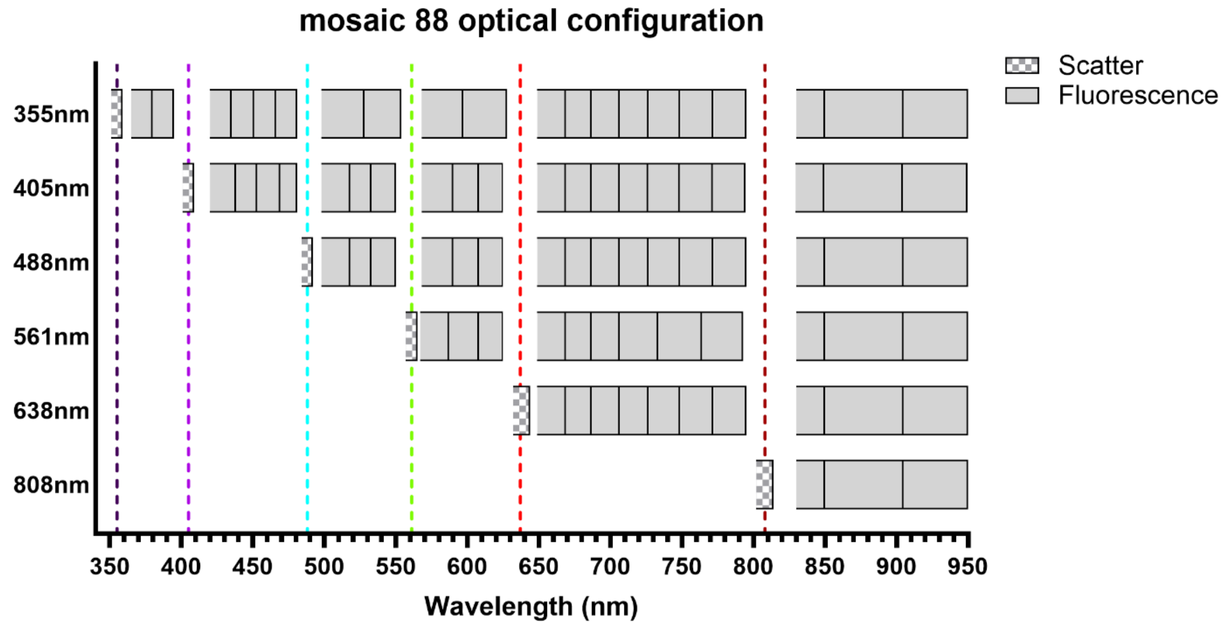


**Supplemental figure 1. Mosaic 88 optical configuration.** The mosaic 88 Spectral Detection Module is equipped with 6 lasers, each with its own SSC detector and 3-20 detectors for fluorescence. Blocks indicate individual detectors; dashed lines indicate lasers.

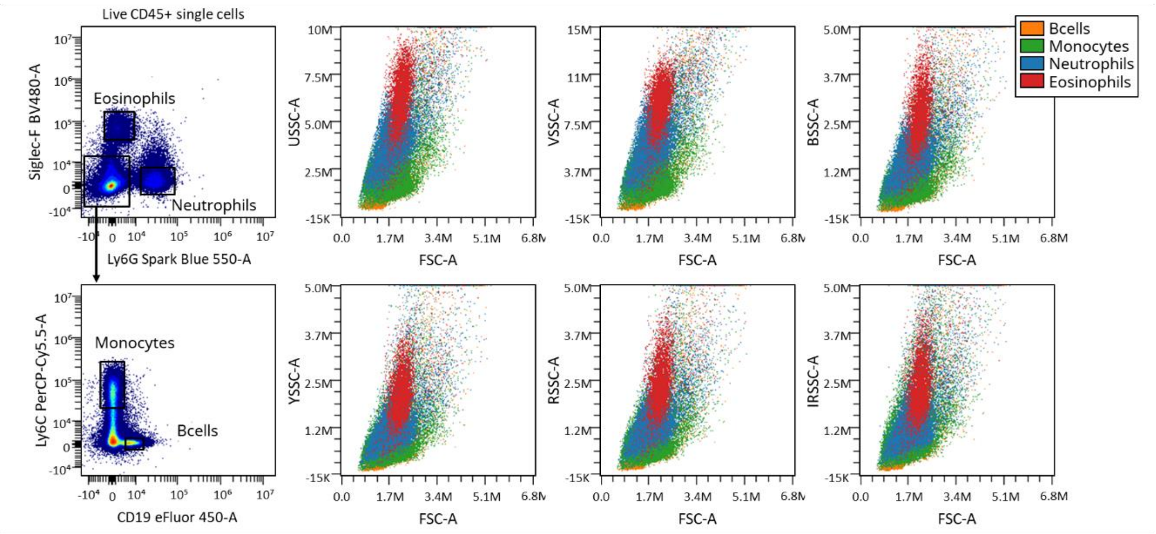


**Supplemental figure 2.Murine eosinophil IRSSC.** Cells were gated as shown on the left. Plotting gated populations as SSC versus FSC for all SSCs showed that murine eosinophils cannot not be distinguished from other cell populations using IRSSC.

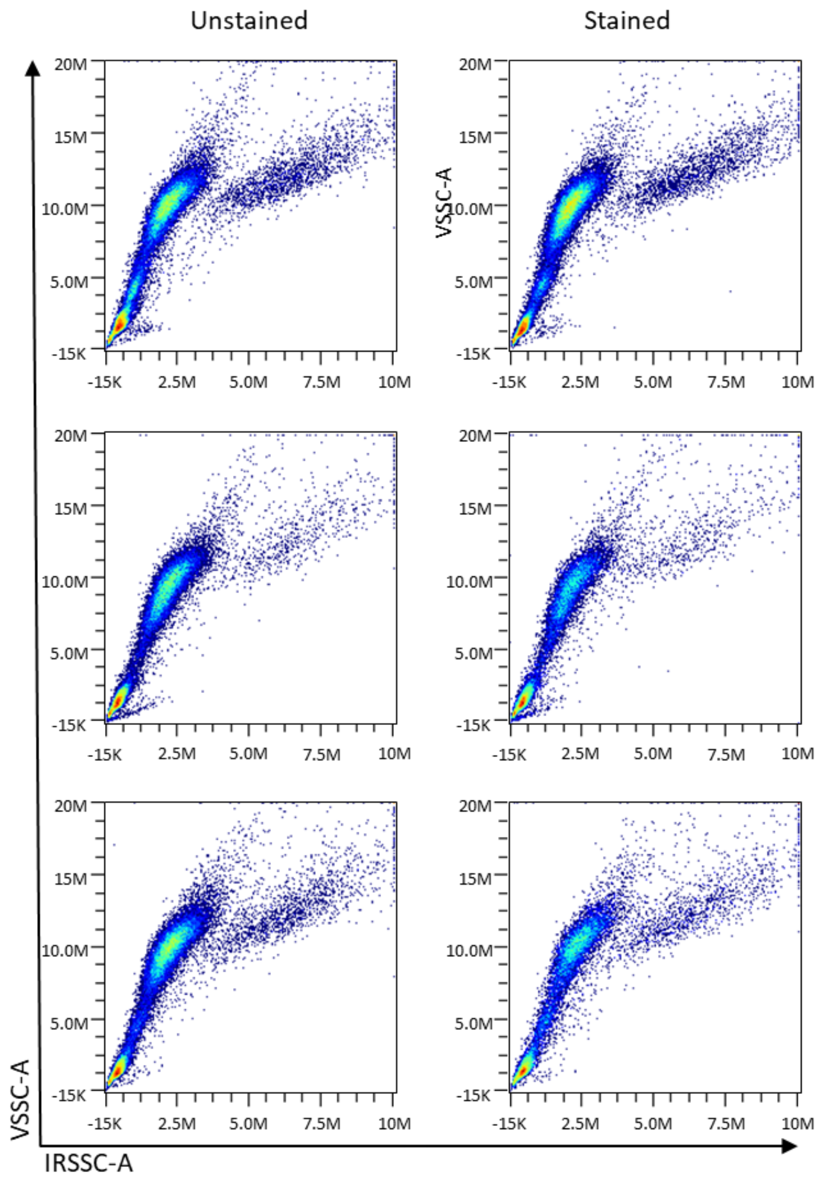


**Supplemental Figure 3. High eosinophil IRSSC does not depend on antibody staining.** Samples stained with antibodies (right) or without antibody staining (left) all showed a similar pattern of IRSSC$^{high}$ cells. Three representative samples from donors with a range of eosinophil frequencies.
